# The potential role of liquid-liquid phase separation in the cellular fate of the compartments for unconventional protein secretion

**DOI:** 10.1101/2023.12.19.572348

**Authors:** Luis Felipe S. Mendes, Carolina O. Gimenes, Marília D. O. da Silva, Saroj K. Rout, Roland Riek, Antonio J. Costa-Filho

**Affiliations:** São Carlos Institute of Physics, University of São Paulo, Avenida João Dagnone 1100, São Carlos, SP 13563-120, Brazil; Molecular Biophysics Laboratory, Department of Physics, University of São Paulo, Ribeirão Preto, SP, Brazil; Laboratory of Physical Chemistry, Swiss Federal Institute of Technology, ETH Hönggerberg, Vladimir-Prelog-Weg 2, 8093 Zürich, Switzerland

## Abstract

Eukaryotic cells have developed sophisticated mechanisms for biomolecule transport, particularly under stress conditions. This study focuses on the unconventional protein secretion (UPS) pathways activated during starvation stress, facilitating the export of soluble leaderless proteins. Specifically, we examine the role of the GRASP family, which is crucial in these processes but has largely unexplored mechanisms, especially regarding the biogenesis and cargo recruitment for the vesicular-like compartment for UPS. Our results reveal that under the circumstances mimicking starvation conditions in yeast, liquid-liquid phase separation (LLPS) is instrumental in the self-association of Grh1, the yeast homologue of GRASP. This association is likely a precursor to the biogenesis of the so-called Compartment for Unconventional Protein Secretion. We demonstrate that Grh1’s self-association involves complex interactions and is regulated by electrostatic, hydrophobic, and hydrogen-bonding forces. Importantly, our study shows that phase-separated states of Grh1 can recruit UPS cargo under starvation-like situations, a significant finding given the minimal interaction between these proteins under normal conditions. Additionally, we address how droplet ageing might affect the ability of cells to return to normal post-stress states. Our findings can potentially enhance our understanding of intracellular protein dynamics and their adaptive responses to stress in eukaryotic cells.

## Introduction

Protein transport within the secretory pathway involves multiple organelles, including the endoplasmic reticulum (ER) and the Golgi complex. Estimates suggest that up to 36% of the total proteome in *Homo sapiens* is predicted to enter this conventional secretory system [1]. This pathway is pivotal for eukaryotic cells as it facilitates the journey of a nascent protein from the ER through the Golgi complex and, ultimately, to its final destination. Despite its significant importance, the ER-Golgi-Final Destination pathway is not the only route for protein secretion within the cell [2]. An increasing number of other proteins is shown to utilize pathways that do not necessitate most conventional machinery components [2,3]. Although unique in their characteristics, those pathways of unconventional protein secretion (UPS) share a commonality in their stress-related triggers [2], including nutrient starvation [4], temperature [5], and ER stress [6]. They handle different protein types, ranging from those membrane-associated translated into the ER and bypassing the Golgi complex to reach their final destinations (Type IV UPS) to soluble proteins in the cytosol recruited for additional functionalities and without ER and Golgi participation (Type III UPS) [2]. Other proteins are directly transported across the membrane by specific ABC transporters (Type II UPS) or through pore-mediated translocation (Type I UPS). This diversity of pathways underlines the astounding complexity of protein transport in eukaryotic cells. The role of UPS is undeniably evident in processes like inflammation, cancer cell proliferation, and the secretion of virulence factors in parasites [7,8], thus amplifying the attention and interest towards UPS to fulfill the continuing gaps in understanding most molecular triggers and cargo pathways.

Types III and IV UPS pathways imply secretion processes that include vesicular intermediates such as autophagosomes, endosomes, multivesicular bodies, and lysosomes [8]. These pathways have been observed across multiple eukaryotes, spanning plants, yeast, flies, and mammals [2,8]. They are directly involved in the export of various protein families, including cytokines, lipid chaperones, hydrolytic enzymes, and aggregate-prone toxic proteins, among others [2,9–11]. A common feature in Types III and IV UPS and the classical secretory pathway is the critical participation of a family of Golgi-matrix proteins, the Golgi Reassembly and Stacking Proteins (GRASPs) [12–16]. GRASPs are evolutionarily conserved peripheral membrane-associated proteins, first identified for their role in reassembling Golgi cisternae following mitotic events [17,18]. They constitute a group of versatile proteins distinguished by unique structural properties [12,13,19–22], several different functionalities [13,23], and a broad interactome [21]. Under cellular stress, GRASPs are translocated from the Golgi to play a central role in UPS. Using the budding yeast *Saccharomyces cerevisiae* as a model system, it was demonstrated that the GRASP65-homologue protein 1 (from now on referred to as Grh1) is incorporated into what is believed to be membrane-like structures adjacent to Sec13-containing ER exit sites [24]. Named the Compartment for Unconventional Protein Secretion (CUPS), this structure was identified as part of the Type III UPS pathway [24]. Starvation induced the maturation of CUPS from immature forms to stable structures, which contained the secretory cargo and were later enclosed by a membrane-bound saccule with a heterogeneous population of vesicles and tubules within [25]. The formation of CUPS was stress-triggered, leading to the substantial accumulation of Grh1 at specific cellular sites [24,26]. This accumulation was fully reversible upon alleviating stress conditions [24]. CUPS formation and dissociation did not rely on new protein synthesis [24,25,27]. Despite these insights, significant unknowns remain, including the molecular mechanisms underpinning CUPS formation, the regulation of its reversibility, and how it selectively recruits UPS cargo.

The self-association of Grh1 into amyloid-like aggregates during Type III UPS and their intracellular colocalisation with CUPS has been previously suggested [26]. However, the regulatory mechanisms guiding their role in UPS remain elusive. Also unclear is what governs the evolution of the CUPS under yeast starvation, especially from a state where Grh1 punctual localization inside these intracellular compartments ceases to be reversible. Given that proteins in physiological conditions stay near their supersaturation concentrations [28], many can sample the three most prevalent states: native, droplet, and amyloid [29]. Eukaryotic cells frequently utilise the droplet state to organise their intracellular components into what are known as membrane-less organelles [29–31]. Ever since the initial observation of this phenomenon in P-granules [32], a multitude of membrane-less organelles have been identified, including but not limited to nucleoli, Cajal bodies, DNA damage foci, the X-chromosome inactivation (XCI) centre, paraspeckles, and stress granules [29]. The unique arrangement of these biomolecules, brought about through liquid-liquid phase separation (LLPS), equips the cell with an exceptional and highly adjustable organisational tool. Beyond their mere organisational utility, membrane-less organelles can modulate a range of biochemical reactions and boost cellular fitness during periods of stress [33–35].

In this study, we observed for the first time that the Grh1 protein forms condensate through LLPS under conditions that simulate starvation in *S. cerevisiae* yeast cells. Our findings highlight that the behaviour of Grh1 is markedly affected by various environmental factors, such as pH levels, membrane surface, and physicochemical stress. Under prolonged incubation, this protein exhibits a dual nature: it can either develop amyloid-like fibres or undergo LLPS, with a specific response depending on the cellular environment. Further, our research indicates that these liquid condensates play a crucial role in attracting UPS cargo, hinting at a potential mechanism for cargo selection in cells. Our study provides significant insights into how proteins like Grh1 adapt their behaviour in response to different stressors and conditions. It offers a more detailed view of the complex protein dynamics within cells under various physiological states.

## Results and discussions

### Grh1 forms amyloid-like fibres in neutral but not in starvation pH

UPS pathways are activated in response to cellular stress, suggesting a mechanism whereby cells navigate stress conditions until homeostasis is re-established. Under nutrient starvation, the budding yeast *S. cerevisiae* provides the best-studied model for activating Type III UPS pathways [10,24]. Nutrient starvation elicits various intracellular physicochemical changes, with a substantial pH decrease being the most notable [36]. While the cytosolic pH under growth conditions is around 7.2-7.4, it can drop to a highly acidic environment of pH 5.4 during glucose starvation [36]. Previous observations have shown that in *S. cerevisiae* cells under starvation stress, Grh1 develops amyloid-like signatures following extended incubation periods, during which the assembly of CUPS transitions from reversible to irreversible [26]. Since protein aggregation is primarily influenced by protein solubility and pH is a potent instigator of protein aggregation, we began by examining the biophysical properties of Grh1 under pH 7.4 for normal conditions (NC) and pH 5.4, mimicking starvation conditions (SC). We used far- and near-UV Circular Dichroism (CD) and size-exclusion chromatography to determine that Grh1 maintained its monomeric state and overall secondary and tertiary structures at room temperature under both conditions (Figure 1A and S1).

**Figure 1:**
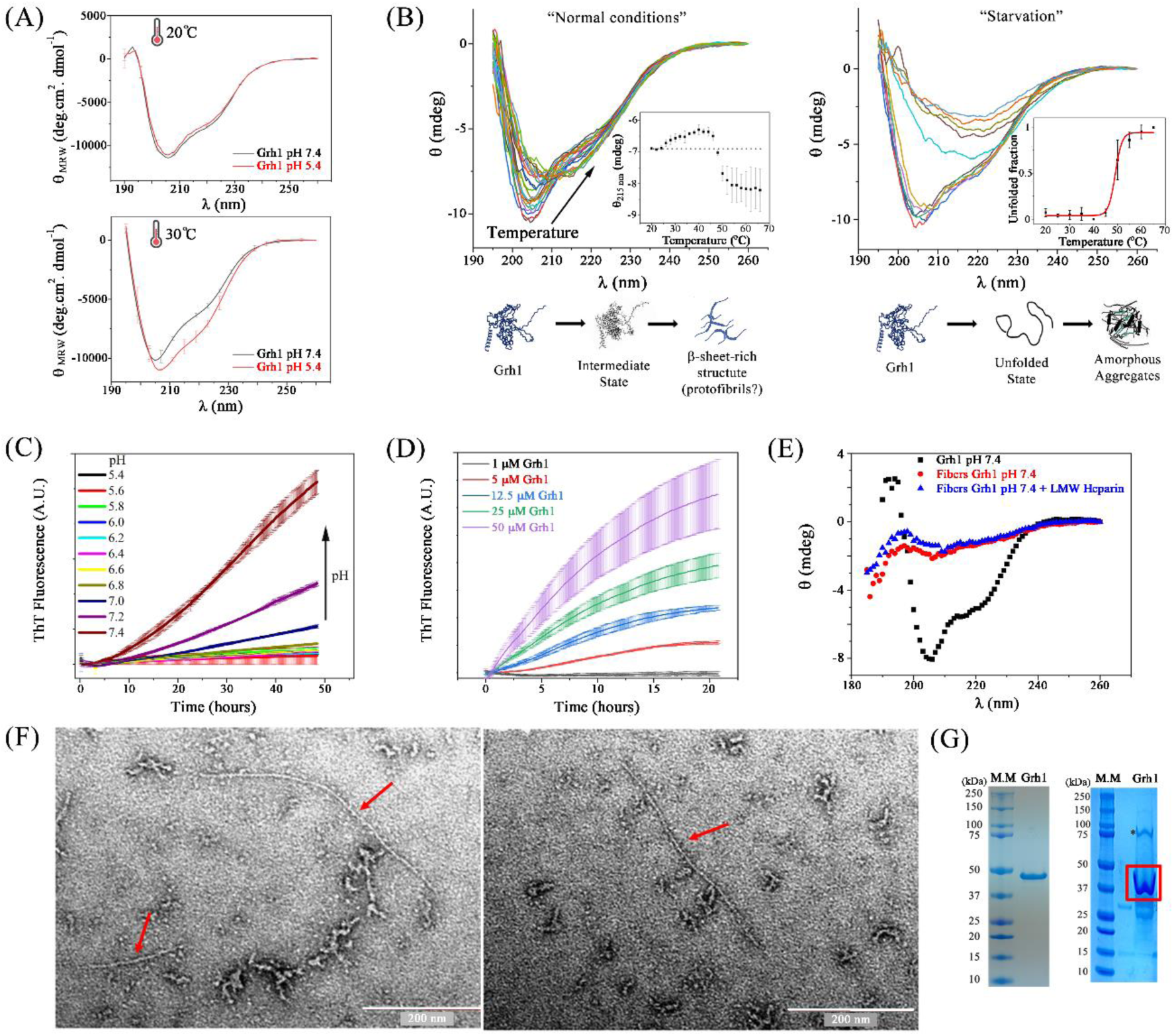
Amyloid-like aggregation of Grh1 studied using spectroscopies and electron microscopy. (A) Far-UV CD spectra of Grh1 in the relevant temperatures of 20 and 30℃. (B) Thermal denaturation of Grh1 (0.1 mg/mL) in starvation and NC monitored by far UV CD. (C) Amyloid-like aggregation was monitored by ThT fluorescence under a wide range of pHs covering starvation (5.4) up to NC (7.4). The McIlvaine buffer solutions were used. (D) Amyloid-like aggregation as a function of Grh1 concentration under NC. (E) Far-UV CD spectra of Grh1 (0.1 mg/mL) and its amyloid-like forms under NC and with the addition of LMW Heparin after 2 days of incubation at 30℃. (F) Representative negative-staining Electron Microscopy micrographs of amyloid-like structures and protofibrils formed by Grh1 at pH 7.4. (G) SDS-PAGE of purified Grh1 (left) and SDS-resistant amyloid-like aggregates of Grh1 observed using SDS-PAGE (right).

Grh1 thermal unfolding at NC presented an intermediate state around 30℃ before its completion with a T_M_ of 45℃ (Figure 1B). Intriguingly, the protein formed soluble β-rich structures at the end of the unfolding experiments (Figure 1B). This pattern was not observed under starvation conditions, where the protein followed a sigmoidal irreversible transition (inset in Figure 1B), culminating in the formation of large, insoluble, and amorphous aggregates at high temperatures (Figure S2). Elevating the temperature to 30℃, optimal for *S. cerevisiae* growth, revealed a partial unfolding of Grh1 in NC that was not mirrored in SC (Figure 1A). This partial unfolding in NC increased the amyloid propensity of Grh1, as suggested by our ThT-binding assay (Figure 1C). Such a tendency was notably mitigated as pH decreased to starvation conditions (Figure 1C). The kinetics of Grh1 aggregation under NC adhered to a classical sigmoid-like pattern, with a strong dependence on protein concentration (Figure 1D), and culminated in a product enriched in β-sheet structures (Figure 1E). Examining Grh1 via electron microscopy under NC revealed the formation of amyloid- and protofibril-like structures (Figure 1F), exhibiting substantial resistance to SDS (Figure 1G). Low molecular mass carbohydrates and glycosaminoglycans are highly concentrated in the Golgi apparatus and secretory vesicles [37]. Therefore, we also tested the influence of low molecular weight heparin in the Grh1 amyloid aggregation, but we could detect significant changes in the amyloid kinetics (Figure 2A).

**Figure 2:**
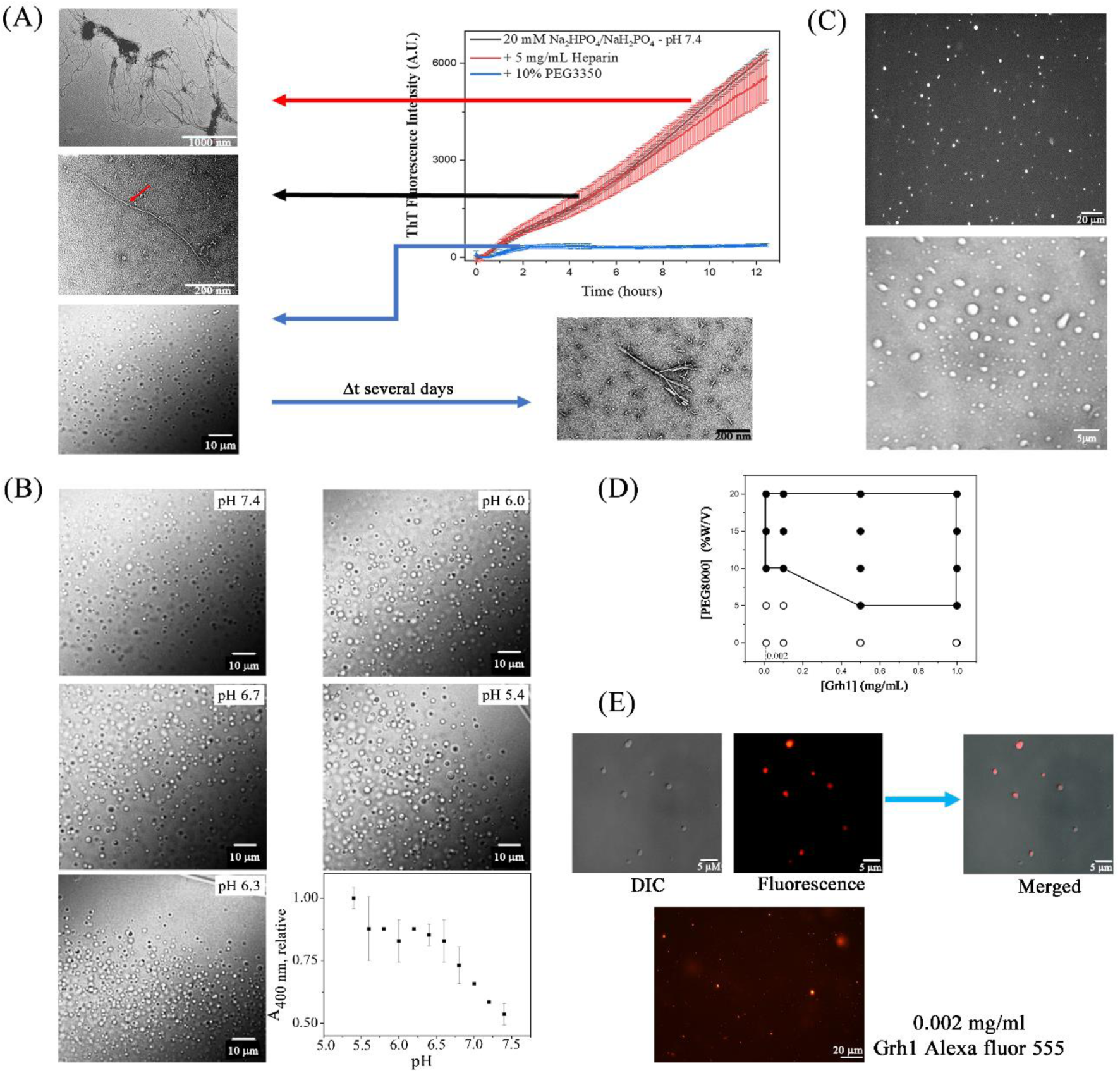
Characterization of Grh1 Liquid-Liquid Phase Separation in Crowded Environments. (A) Monitoring Grh1’s amyloid-like formation using Thioflavin T (ThT) fluorescence. The aggregation process of Grh1 can be modulated by heparin. However, the timescale for amyloid-like aggregation can be significantly reduced only through LLPS. Electron microscopy images were obtained to visualize amyloid-like structures, while Differential Interference Contrast (DIC) microscopy was used to identify droplets formed due to LLPS. (B) The liquid-like characteristics of Grh1 droplets formed under starvation were validated by the rapid permeation of the small fluorescent probe Sypro-Orange (upper panel) and their wetting properties on microscope glass slides (lower panel). (C) Representative DIC microscopy images of Grh1 droplets under a pH range from 7.4 to 5.4. Images were captured focusing on the interfacial region between the edges of the solution and the air. (D) The phase diagram of Grh1 under starvation conditions was constructed using DIC microscopy. Black points represent conditions where liquid droplets were observed. The median Grh1 abundance in yeast cells is 4576 ± 1041 molecules per cell, corresponding to a protein concentration ranging from 100 nM to 1µM (∼0.04 – 0.004 mg/mL). A fluorescence microscopy image of Alexa-Fluor 555-labelled Grh1 was obtained at a protein concentration of 0.002 mg/mL under starvation conditions.

Although Grh1 has been previously observed to form amyloid structures under harsh *in vitro* conditions at temperatures surpassing its T_M_ [5], this propensity is also evident at physiologically relevant temperatures under various native conditions, not starvation. Moreover, the observations presented here contrast with previous findings *in cells*, which indicated that the amyloid signature for Grh1 manifested exclusively in cells experiencing starvation, and this was associated with the formation of CUPS during extended periods of stress [26]. Therefore, we sought strategies to reconcile those previous observations with our current *in vitro* analyses.

### Grh1 shows a high tendency for liquid-liquid phase separation (LLPS)

The liquid droplet state formed through LLPS shows high conformational entropy, maintained by many feeble intra- and inter-molecular interactions [38,39]. Thus, a strong correlation between intrinsic disorder – a trait often found in polyvalent polymeric biomolecules – and *in vivo* condensation events has been demonstrated [39,40]. Grh1 is a protein enriched in intrinsically disordered and amyloidogenic regions (Figure S3), which have not yet been checked for their propensity to undergo LLPS. To test for this, we first used several predictive algorithms that indicated Grh1 exhibited a pronounced tendency for droplet formation via LLPS (Figures S3 and S4). We further examined and experimentally confirmed the potential of Grh1 to form condensates within crowded environments (see Figure 2A and Movie S1). Interestingly, the occurrence of the droplet state considerably lessened the propensity for amyloid aggregation in Grh1 (from hours to several days), as evidenced by ThT fluorescence measurements (Figure 2A).

We next focused on how Grh1’s LLPS changed upon pH variation. Grh1 manifested a pronounced tendency to form liquid-like droplets across an extensive pH range from NC to SC (Figure 2B). Notably, upon pH reduction, we observed a significant increase in the population of droplet states, as tracked by optical dispersion (Figure 2B). These condensates exhibited the hallmarks of liquid droplets: high permeability to small molecules (upper panel in Figure 2C), fusion propensity (Figure S5 and Movie S2), and surface wetting (lower panel in Figure 2C). Additionally, the formation of droplets was independent of the type of macromolecular crowding agent (Figure S6) and the corresponding increased viscosity (Figure S6).

To determine what kind of interactions (electrostatic, hydrophobic, π–π, or cation-π interactions) were mediating the Grh1 phase separation, we examined the effect of increasing the ionic strength (NaCl) and either adding 1,6-hexanediol (aliphatic alcohol known to break weak hydrophobic interactions) or low urea concentrations [41] (Figure S6). Under SC, we observed that Grh1 droplet assembly was mediated by a range of weak interactions, including ionic, hydrophobic, and hydrogen bonds (Figure S6). The low saturation concentration of the protein further substantiates Grh1’s high propensity for droplet formation under starvation-like conditions (Figure 2D). Interestingly, we identified liquid droplets at lower protein concentrations than those typically found *in vivo* for Grh1 (Figure 2D) [42].

### LLPS protects Grh1 from fibrillation in NC

Taking condensates formed via LLPS as a typical organization of Grh1 in crowded environments, we replicated the amyloid kinetic studies of Figure 1D, increasing the Grh1 fraction within the droplet state (by increasing the molecular crowding) while maintaining the previously used parameters. Intriguingly, the droplet state shielded Grh1 from forming amyloid structures under NC (Figure 2A). This propensity towards amyloid formation significantly declined as the protein’s presence within liquid droplets increased (Figure 3A). The formation of droplets acted as a protective mechanism, preventing Grh1 from transitioning into an intermediate folding state under NC, as seen in the thermal denaturation followed by CD at 215 nm (Figure 3B). LLPS possibly shielded the protein’s amyloidogenic regions from exposure.

**Figure 3:**
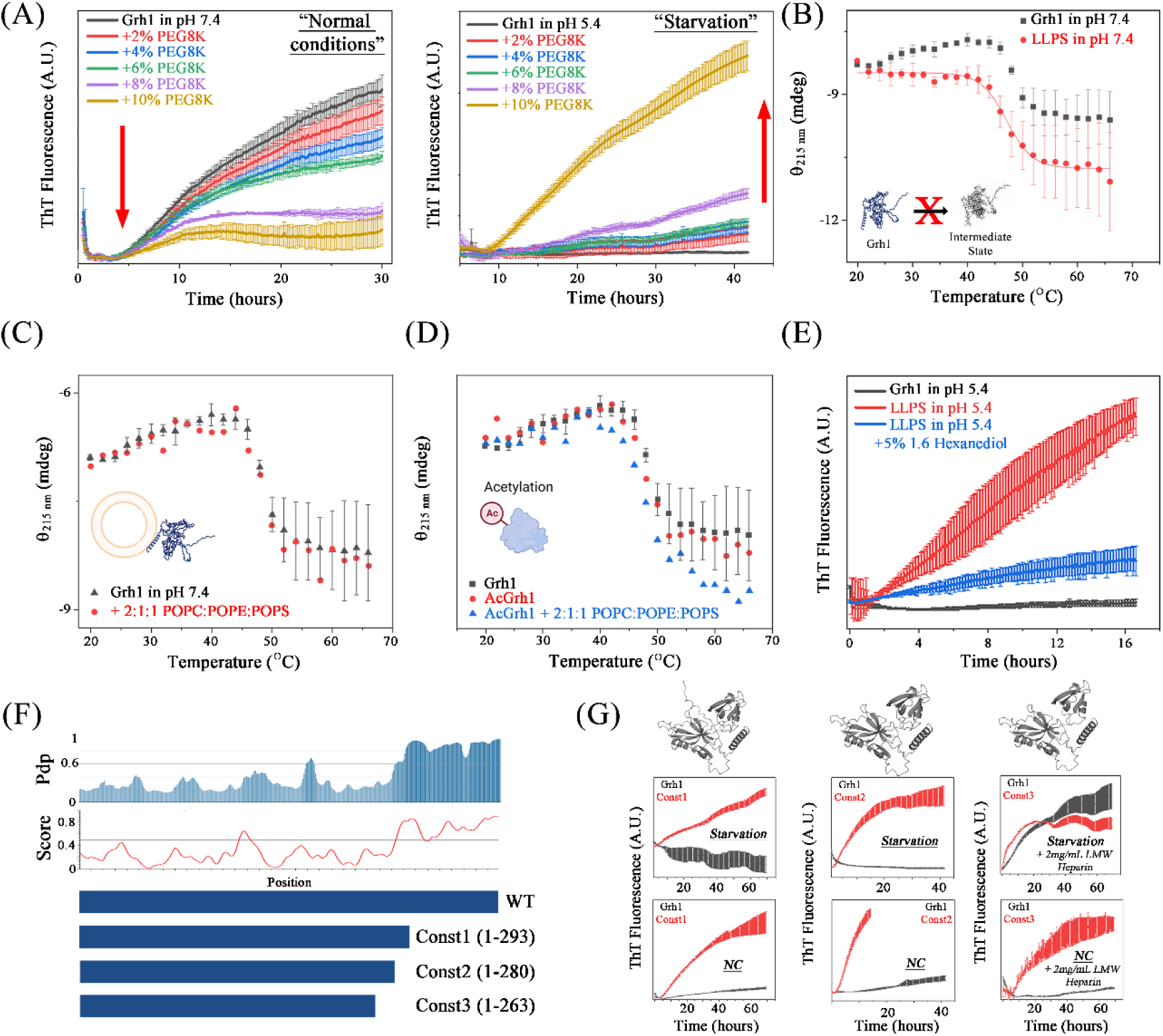
Investigation of Grh1 Liquid Droplet Aging and Influence of Environmental Factors. (A) Kinetics of ThT fluorescence in the presence of Grh1 in a crowded environment under normal (NC) and starvation-like (SC) conditions. The crowding effect was modulated by different concentrations of PEG8000, with the direction of increase in PEG8000 concentration indicated by red arrows. (B) Thermal denaturation of Grh1 in solution and phase-separated states tracked by circular dichroism. (C) Effect of a model membrane simulating the native membrane of *S. cerevisiae* [43] on the formation of the unfolding intermediate state of Grh1, observed through thermal denaturation tracked by circular dichroism. (D) Similar to those in (C), but performed with N-terminus acetylated Grh1 compared to the native protein. (E) Monitoring of amyloid-like aggregation of Grh1 under phase-separated conditions using ThT fluorescence. 1,6 Hexanediol disrupts the weak hydrophobic interactions, which are primarily responsible for the assembly of liquid droplets. (F) Graphical representation of three distinct constructs of Grh1 utilized to progressively remove portions of the intrinsically disordered C-terminus region. The sizes represented by the blue bars correlate with the predicted propensity for LLPS (Pdp – from FuzDrop [44]) and intrinsic disorder prediction (Score – from VSL2 [45]). (G) Kinetics of amyloid-like aggregation of different Grh1 constructs monitored by ThT fluorescence under NC and SC. An increase in ThT fluorescence for construct 3 in conditions without low molecular weight (LMW) heparin was not detected.

Our following action was to examine whether the N-terminal acetylation of Grh1 by the NatC complex - a topic we will elaborate on later - would also halt the formation of Grh1 amyloid structures under NC. Acetylated Grh1 (AcGrh1) preserved the LLPS tendency observed in the non-acetylated version (see Figure S7). It neither impacted the formation of the intermediate state at physiological temperatures (Figure 3D) nor the amyloid-like aggregation under NC (Figure S8). We then explored whether the presence of membrane structures would influence the aggregation of Grh1 and AcGrh1. Even though it slightly altered their melting temperatures (Figures 3C and 3D), the presence of membrane models did not impede protein aggregation, irrespective of a broad array of lipid compositions, ranging from the simplest (pure POPC) to those more akin to natural structures (*S. cerevisiae* Total Lipid Extract). Therefore, the protection against Grh1 amyloid formation under NC could only be achieved through the phase-separated state. Even though this protein possesses an intermediate state with a high propensity to form amyloid-like structures under NC, LLPS shields Grh1 from this transition. This observation might explain why Grh1 does not form amyloid structures *in vivo* despite being close to its solubility limit.

### LLPS is essential for Grh1 fibrillation in SC

The outcomes observed at starvation pH contrast those found under normal conditions. A decrease in pH shifts the equilibrium, favouring the condensed phase (refer to Figure 2B). However, under starvation conditions, Grh1 droplets exhibited an accelerated maturation towards an amyloid-like structure, as tracked by ThT fluorescence (Figure 3A). Subsequently, we scrutinized the droplet state’s response to 1,6-hexanediol. Prior studies have established that 1,6-hexanediol can destabilise the droplet state when hydrophobic interactions contribute to its stability [29]. While the droplet state of Grh1 is not solely or predominantly maintained by hydrophobic interactions (see Figure S5), their perturbation proved sufficient to underscore the liquid-like condensate’s crucial role in the amyloid aggregation since the introduction of 1,6-hexanediol notably attenuated it (Figure 3E). Consequently, we demonstrated that droplet formation via LLPS dually influences Grh1’s fate, contingent upon the cellular conditions.

According to LLPS predictors, the disordered C-terminal region (SPR domain) of Grh1 significantly drives its phase separation into liquid droplets (Figure 3F). Therefore, our subsequent effort sought to assess the impact of this vastly disordered region on Grh1’s amyloid propensity under normal and starvation conditions. We tested three distinct Grh1 constructions to evaluate the SPR influence on droplet formation and the prevalence of regions with a high tendency for amyloid formation (Figure 3F). Intriguingly, by excising the SPR region to varying extents, we noted a dramatic surge in the exponential phase steepness of the kinetic curves in normal conditions. Conversely, the lag phase was considerably shortened (Figure 3G). We identified an amyloid aggregation kinetic curve for constructs 1 and 2, even under starvation conditions (Figure 3G). Thus, our data proved that the disordered region, comprising roughly 25% of Grh1, safeguarded the protein from amyloid aggregation under normal conditions and crowded environments.

The aggregation curves were more significantly impacted for construct 3 (Figure 3G). The key distinction between constructs 3 and 1 or 2 is that, in construct 3, the disordered tail is completely removed along with the final stretch of the PDZ2 domain. We could not detect any amyloid-like aggregation under starvation conditions - a propensity that could only be restored using heparin (Figure 3G). Even under normal conditions, construct 3 only formed amyloid structures in the presence of heparin. Given the identification of amyloidogenic regions localized to the terminal segment of PDZ2 (Figure S9), it is plausible to propose that this region is implicated in the amyloid-like aggregation of Grh1. The adjacency of this segment to the intrinsically disordered SPR domain may facilitate the regulatory mechanism of Grh1 in modulating aggregation via liquid-liquid phase separation.

The disordered SPR domain appears instrumental in regulating LLPS tendency, essential for the protein’s effective response to *S. cerevisiae* starvation conditions. The theoretical pI of the disordered domain is approximately 6.5 (Figure S9), precisely within the range of pH levels that govern normal and starvation conditions. This tendency is not followed by the other regions of Grh1 (Figure S8). Moreover, our findings suggest that the PDZ2 domain exclusively controls the aggregation propensity, and its final portion plays a pivotal role in Grh1’s high aggregation propensity.

### CUPS as a Grh1 enriched phase-separated condensate?

Given that Grh1 liquid condensation is significantly enhanced in response to acidification and most unconventional secretion events are not constitutive but stress-induced [2], we proposed that the CUPS could represent a phase-separated structure formed by a Grh1-rich phase capable of recruiting UPS cargo. Accordingly, the detected amyloid signature previously observed *in cell* [26] might be attributable to droplet ageing, a property observed previously in other well-known amyloidogenic proteins [46–48]. We showed here that Grh1 LLPS is fully reversible, enhanced under starvation conditions, and notably diminished under normal conditions, which are observations aligning with the CUPS dynamics observed *in vivo* [24].

Two other experimental observations need to be validated to strengthen our hypothesis. Firstly, the Grh1 liquid condensate must demonstrate the capacity to recruit UPS cargo under starvation conditions. Secondly, the amyloid signature should be observable as time passes in *in vivo* experiments [26], typically several hours. To address the first point, we used the leaderless protein Acyl-CoA Binding Protein 1 (ACBP, also known as Acp1) as a UPS-cargo model. Due to nutrient or nitrogen deprivation, stress stimulates the unconventional secretion of ACBP in many systems, from *Dictyostelium discoideum* to *S. cerevisiae* [4,49,50]. ACBP lacks a signal peptide and plays a crucial role as an intracellular carrier of acyl-CoA esters [51]. It was shown *in vivo* that ACBP is contained in CUPS and colocalized with Grh1 in *S. cerevisiae* under starvation [25]. Grh1 is fundamental for the UPS of ACBP in all studied models [10].

Under non-phase separated conditions, our spectroscopy measurements failed to detect any binding between Grh1 and ACBP, suggesting that the dissociation constant, if any binding event exists, would likely exceed mM levels (Figure S10). Under our phase-separated conditions, we could not detect any visible ACBP droplet formation (Figure 4). To test the potential ACBP recruitment, we employed fluorescence microscopy under LLPS conditions where Grh1 and ACBP were labelled with Alexa Fluor 555 and fluorescein isothiocyanate (ACBP:Grh1 ratio of 1:1), respectively. ACBP was clearly seen to accumulate within the Grh1 droplets (Figure 4A), indicating that the phase-separated state of Grh1 could recruit UPS cargo under starvation-like conditions.

**Figure 4:**
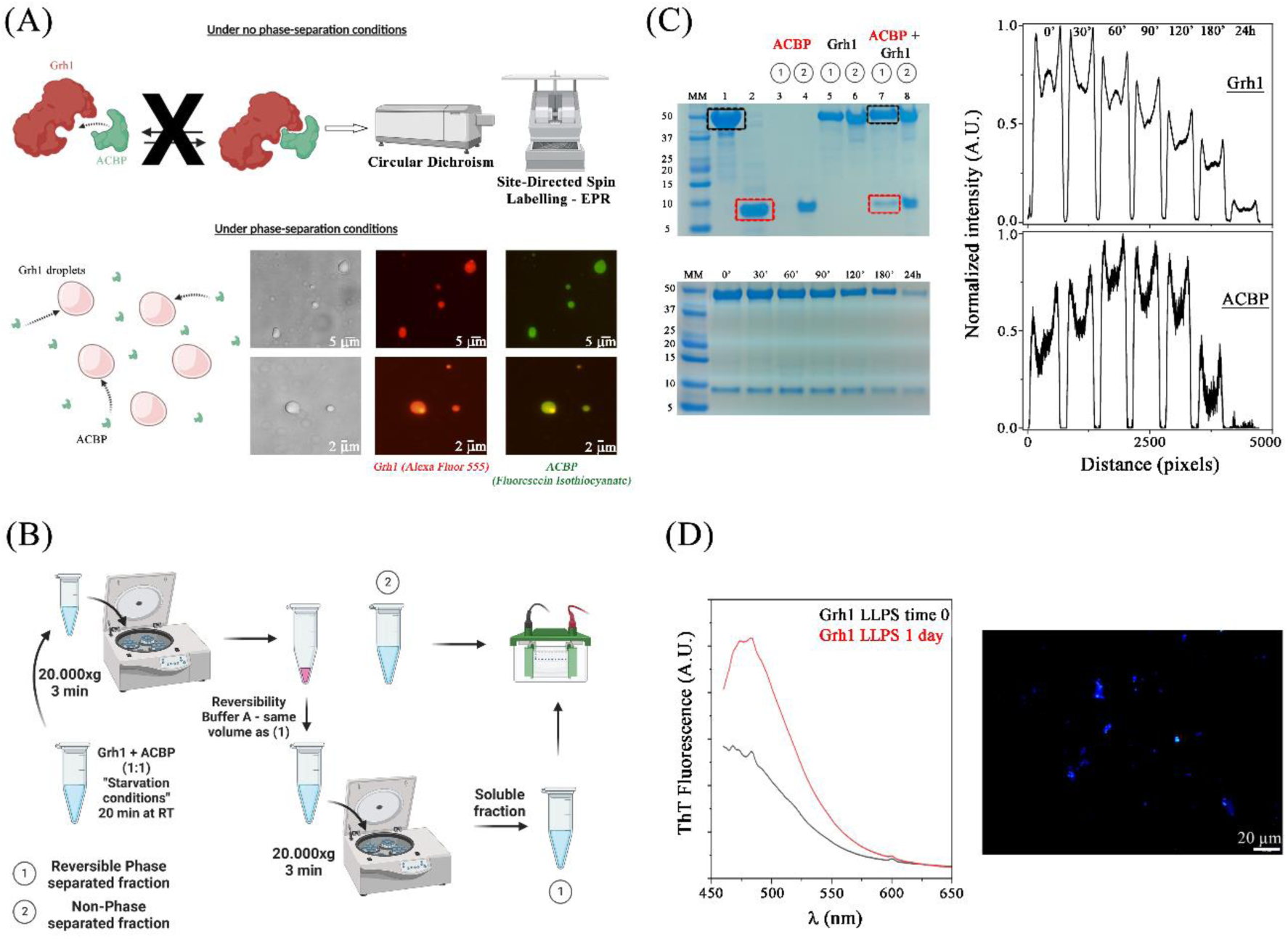
Characterization of the Phase-Separated State of Grh1 in the presence of ACBP. (A) Illustration of ACBP binding to soluble Grh1 and phase-separated states. Direct binding of ACBP to free Grh1 was assessed through circular dichroism and electron paramagnetic resonance using Grh1 labelled with S-(1-oxyl-2,2,5,5-tetramethyl-2,5-dihydro-1H-pyrrol-3-yl)methyl methanesulfonothioate. No binding was detected under the tested conditions, indicating that any possible interaction likely has a K_d_>1mM. Grh1 droplets were imaged via DIC microscopy, and the colocalisation of ACBP into Grh1 liquid droplets was captured through fluorescence microscopy. The colour channels correspond to Alexa Fluor 555-Grh1 (red) and fluorescein isothiocyanate-ACBP (green). Two representative sets of images are shown. (B) Schematic of the protocol designed to separate the (1) reversible phase-separated droplets and the (2) non-phase-separated protein in solution. (C) SDS-PAGE data of samples processed according to the protocol depicted in (B). Grh1 and ACBP bands are indicated by black and red boxes, respectively. The upper gel displays fractions of purified Grh1 (lane 1), ACBP (lane 2), and phase-separated states (as labelled in the image). The numbers 1 and 2 within the circles correspond with the fractions outlined in (B). The lower gel shows the fraction of reversible-phase-separated states as a function of incubation time. The intensity of the bands (Grh1 and ACBP) in the lower gel is represented on the right as a function of pixel distance, extracted from the gel image and using Fiji software. (D) The non-reversible fraction of Grh1+ACBP was analyzed using ThT fluorescence and deep-blue autofluorescence.

In order to ascertain that only reversible Grh1 droplets are instrumental in ACBP recruitment, we designed a protocol to examine recruitment over time. This protocol isolated the phase-separated fraction via centrifugation, followed by solubilization using the purification buffer (where reversibility must be accomplished) and further centrifugation to eliminate precipitation (Figure 4B). This approach verified Grh1’s ability to recruit ACBP and assessed LLPS reversibility over time. The protocol confirmed that ACBP did not form liquid droplets under starvation-like conditions and remained in the soluble phase after the initial centrifugation (Figure 4C). Conversely, Grh1 was observed in both the soluble and reversible phase-separated states, indicating that under the tested conditions, approximately half of the protein remained in the condensed phase. When both proteins were simultaneously present, ACBP began to be found in the phase-separated population formed by Grh1 (Figure 4C).

The fraction of reversible Grh1 droplets diminished over time under starvation conditions (Figure 4C). ACBP’s incorporation within Grh1 droplets intensified over time (up to ∼one hour), coinciding with the period when the reversibility of Grh1 droplets started to be significantly impacted (Figure 4C). The isolated non-reversible structure formed by Grh1 over time under starvation exhibited an amyloid-like signature characterized by an increase in ThT affinity and the development of deep-blue autofluorescence (Figure 4D) [26,52]. These observations agree with the previous *in vivo* data of *S. cerevisiae* under starvation for prolonged times [26]. Liquid protein droplets can inherently concentrate proteins at strikingly high levels. Consequently, this situation creates a unique setting to bypass the high entropically-cost nucleation of amyloid fibril formation. The connection between LLPS and amyloidogenesis has been previously observed in several human diseases, such as Amyotrophic Lateral Sclerosis (ALS), Alzheimer’s Disease (AD), and Parkinson’s Disease (PD) [29,53].

Given that Grh1 exhibits a high propensity to form amyloid-like aggregates in our *in vitro* studies at NC, it is expected that we detected an amyloid signature in Grh1 droplet samples under SC after prolonged incubation (Figure 3A). This observation aligns with the limited tolerance of *S. cerevisiae* to starvation conditions [54] and the irreversible Grh1 localization after prolonged starvation observed *in cell* [24,25]. Burns et al. showed that the Grh1-containing compartment in *S. cerevisiae* under starvation was stable for up to 8 hours [24]. Our kinetic experiments in Figure 3A do not entirely agree with this *in vivo* observation, as the timescales for complete aggregation do not match. Thus, we examined the Grh1 aggregation kinetics’ dependence on various physicochemical parameters, including Hofmeister ions and heparin (Figure 5). The introduction of Low Molecular Weight (LMW) heparin (Figure 5A) and Mg^2+^ ions (Figure 5B) significantly reduced the lag phase to a timescale of about an hour. Therefore, the kinetics of Grh1 aggregation can be substantially modulated by several parameters present in the secretory pathway. The 1-h timescale coincides with *S. cerevisiae*’s response time under starvation, explaining the previously observed *in vivo* amyloid signature [26].

**Figure 5:**
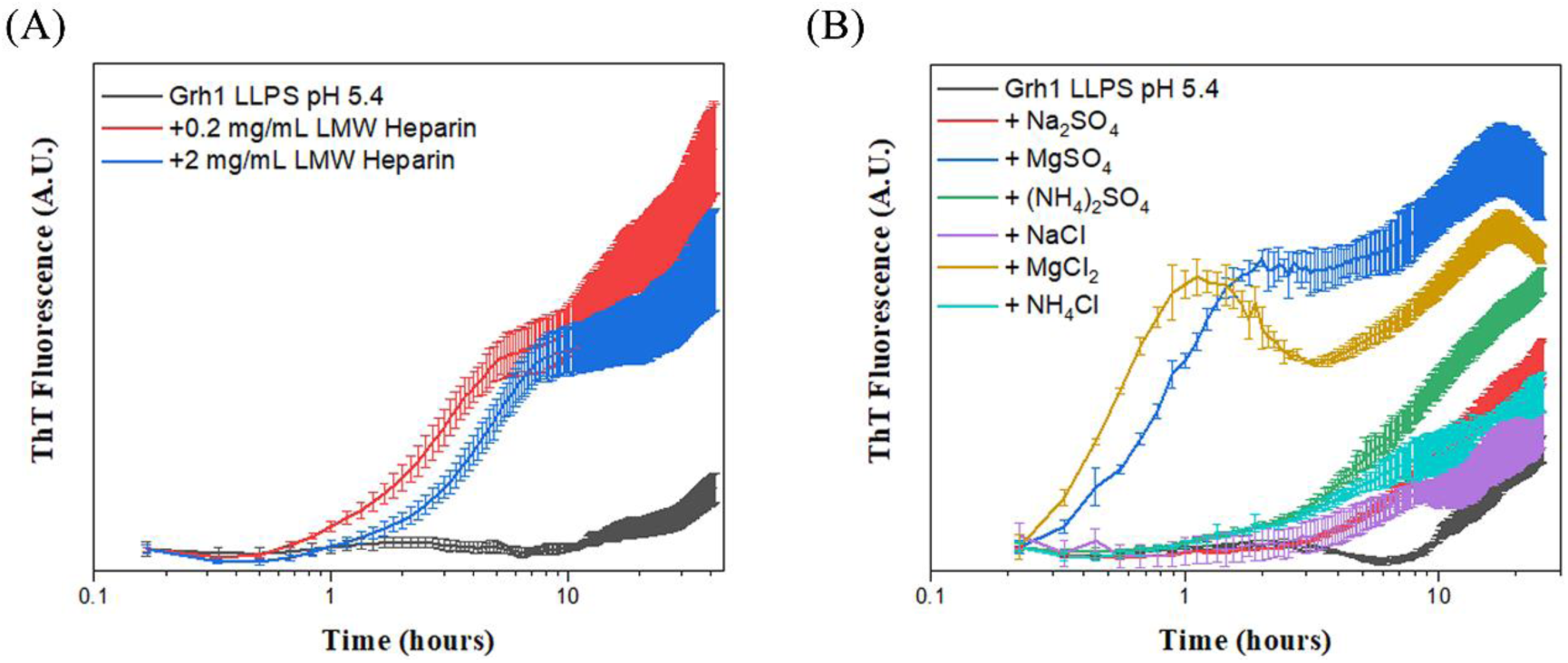
Monitoring Grh1 Liquid Droplet Aging and Transformation into Amyloid-Like Structures via ThT Fluorescence. This figure illustrates the progression of Grh1 liquid droplets into amyloid-like structures, as detected by Thioflavin T (ThT) fluorescence, a diagnostic dye for amyloid structures. The impact of various molecules on the aggregation of Grh1 was evaluated. (A) Influence of Low Molecular Weight (LMW) Heparin on Grh1 Aggregation. (B) Effect of Salts on Grh1 Aggregation. In this analysis, we explored the Hofmeister series. The Hofmeister series is typically arranged regarding the ion’s ability to “salt in” or “salt out” proteins. SO4-is more kosmotropic than Cl-; for cations, this follows as NH ^+^ > Na^+^ > Mg^2+^ from more kosmotropic to more chaotropic.

## Conclusions

The recent findings of LLPS of biomolecules within the intracellular environment, leading to the creation of membrane-less organelles, is heralded as a transformative advancement in cell biology [31,32,55–57]. The multitude of novel and intriguing ways living cells utilize this physicochemical process continues to expand. Here, we have demonstrated that Grh1, a Golgi matrix protein implicated in the Unconventional Protein Secretion (UPS) of various protein cargoes, exhibits a strong propensity for liquid droplet formation via LLPS under functionally relevant conditions. This propensity is amplified under circumstances wherein Grh1 plays a role in the Compartment for Unconventional Protein Secretion (CUPS) biogenesis. Our study reveals that Grh1 liquid droplets can recruit UPS cargo, suggesting a possible mechanism for this biological phenomenon. We propose that CUPS are membrane-less organelles predominantly composed of Grh1. This postulation provides a plausible explanation for the observed reversibility, ageing, and unusual cargo recruitment previously reported in *in vivo* studies [10,24–26].

Additionally, we found that Grh1 has a high inclination to form amyloid-like structures under native conditions. However, we also discovered that LLPS protects Grh1 from transitioning into amyloid structures by precluding the formation of an intermediate state under physiological conditions. These observations exemplify LLPS’s dual role in dictating a protein’s fate under differing cellular conditions (Figure 6). Our findings enrich our understanding of LLPS’s observable participation in biological processes and pave the way for future exploration of UPS mechanisms.

**Figure 6:**
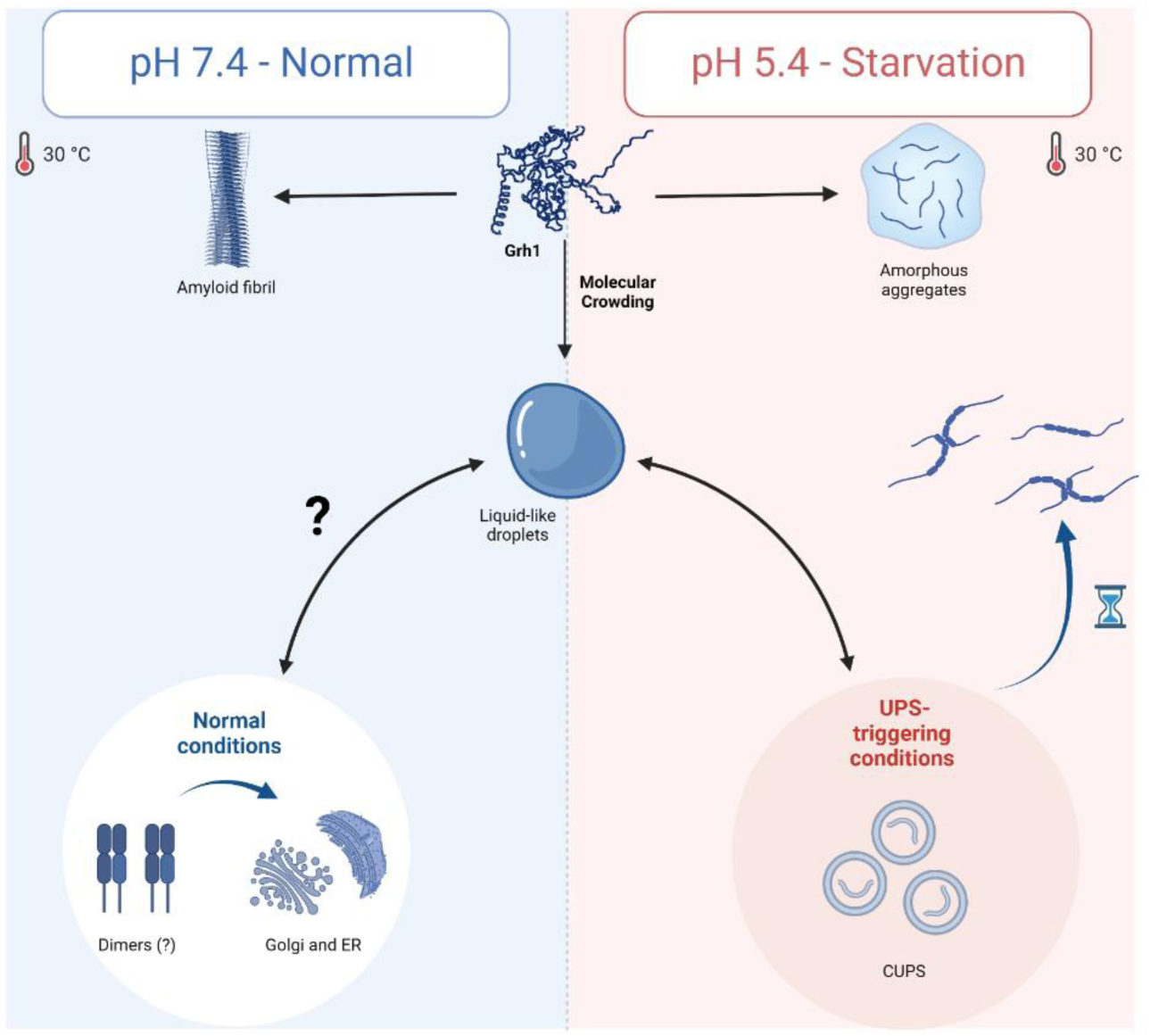
Schematic model of Grh1 fate under different cell/physicochemical conditions observed in normal conditions and starvation.

## Material and methods

### 2.1 Protein Expression and Purification

The yeast Grh1 open reading frame (Uniprot Q04410) was synthesised with codon optimisation for *E. coli* expression by GenScript^®^ Biotech and subcloned in pET-22b (+) vector using the Nde1 and Xho1 restriction sites. Protein expression was performed in Bl21 (DE3) pLys competent cells (Novagen) using an L.B. medium. Cells grew at +37 °C in the presence of 100 mg/L ampicillin. Recombinant gene expression was induced by 0.5 mM IPTG at OD600 = 0.8. Expression was performed at +18 °C for 16 h. Harvested cells were lysed and resuspended in Buffer A (10 mM HEPES/NaOH, 300 mM NaCl, 5 mM 2-MercaptoEthanol, 10% Glycerol, pH 7.4) in the presence of 1 tablet of cOmplete™, EDTA-free Protease Inhibitor Cocktail and 0.5 mg/ml lysozyme under constant stirring for 20 minutes at room temperature. Afterward, cells were disrupted in 110S microfluidiser (Microfluidics, Newton, Massachusetts, U.S.A.) at 80,000 psi. The lysate was centrifuged at 25.000 x g (rotor 45 Ti, Optima L-90K Ultracentrifuge, Beckman Coulter International S.A., Nyon, Switzerland), +4 °C for 25 minutes. The soluble fraction of the disrupted cell solution was loaded into a Ni-NTA column system (Qiagen). The column was washed with a volume corresponding to 5 column volumes (CV) of Buffer A, Buffer A + 10 mM Imidazole, and Buffer A + 15 mM Imidazole. The bound protein was eluted using 3 C.V.s of Buffer A + 500 mM imidazole. The protein solution was concentrated using a 30 kDa cutoff centrifugal filter unit (Millipore, Burlington, MA, U.S.A.) and loaded into a Superdex200 10/300 GL gel filtration column (G.E. Healthcare Life Sciences, Chicago, IL, U.S.A.) coupled to an Äkta purifier system (G.E. Healthcare). The purification efficiency was monitored by electrophoresis using NuPAGE® Novex® 4-12% Bis-Tris Gels (Invitrogen™ by LifeTechnologies™, LuBioScience GmbH, Lucerne, Switzerland) according to the manufacturer’s instructions. Grh1 concentration was determined by U.V. spectroscopy (ε_280 nm_ = 32890 M^−1^.cm^−1^).

The different Grh1 constructions, named construct 1 (1-263), construct 2 (1-280) and construct 3 (1-298), were expressed and purified using the same protocol. Acetylated Grh1 was produced by co-expressing the Grh1 – pET22b vector with the yeast NatC complex (formed by full-length Naa30, Naa35, and Naa38) cloned in pRSFDuet-1 (gently given by prof. O. Daumke [58]). The NatC complex does not carry a HisTag in this strategy and is completely removed during the Ni-NTA purification step.

The yeast Acyl CoA Binding Protein open reading frame (Uniprot P31787) was amplified from the *Saccharomyces cerevisiae* strain ATCC 204508/S288c genome and cloned in the pETSUMO vector using BamH1 and Xho1 restriction sites. ACBP was purified using a similar protocol for Grh1 with changes only after the Ni-NTA purification. Ni-NTA purified ACBP was mixed with ULP1 (1/1 ratio in mass) and dialysed overnight at 4℃ against a reservoir (volume 100 times higher) composed of 20 mM Tris/HCl pH 8.0, 20 mM NaCl and 5 mM 2-Mercaptoethanol. The solution was reloaded in a Ni-NTA column, and the flowthrough fraction (composed of HisTag-free ACBP) was collected. The solution was concentrated using a 3 kDa cutoff centrifugal filter unit (Millipore, Burlington, MA, USA) and loaded into a Mono Q 5/50 GL (GE Healthcare Life Sciences, Chicago, IL, USA) coupled to an Äkta purifier system (GE Healthcare). The protein solution was then submitted to a linear gradient of NaCl from 20 to 500 mM (+ 20 mM Tris/HCl pH 8.0) at a flow rate of 1 mL/minute with a 2 mL injection volume. The peak associated with pure ACBP was concentrated and further purified by SEC using a Superdex 75 (HR 10/30) column coupled to an Äkta purifier system (GE Healthcare) using Buffer A. ACBP concentration was determined by U.V. spectroscopy (ε_280 nm_ = 18.450 M^−1^.cm^−1^).

### 2.2 Membrane Preparation

1-palmitoyl-2-oleoyl-glycero-3-phosphocholine (POPC), 1-palmitoyl-2-oleoyl-sn-glycero-3-phosphoethanolamine (POPE), 1-palmitoyl-2-oleoyl-sn-glycero-3-phospho-L-serine (POPS) and Yeast Total Lipid Extract (*S. cerevisiae*) were purchased from Avanti Polar Lipids Inc. (Alabaster, AL). A dry lipidic film (POPC, POPC/POPE/POPS (2:1:1), or Yeast Total Lipid Extract) was prepared in a round-bottom glass tube from a CHCl_3_ solution by removing the solvent under a constant stream of N_2_ followed by incubation under vacuum for 16 hours to remove traces of organic solvent. Multilamelar vesicles (MLV) were prepared from the dispersion of the lipids in the appropriate buffer, incubated at RT (POPC or POPC/POPE/POPS 2:1:1) or 60℃ (Yeast Total Lipid Extract) followed by 3 rounds of rapid freeze-thawing. Large Unilamellar Vesicles (LUVs) were obtained by subjecting the lipid dispersion to 21 passes through polycarbonate membranes of 100-nm pore size (Nuclepore Corp., Cambridge, CA) using an Avanti Mini Extruder (Alabaster, AL, USA). Small Unilamellar Vesicles (SUVs) were prepared from the lipid dispersion by sonication in ice-cold with a probe microtip sonicator (Vibra-Cell™ - Sonics) for 10 minutes (5-s on and off cycles) at 20% amplitude.

### 2.3 Aggregation Kinetics and Turbidity Measurements Using Microplate Reader

ThT from Sigma (St. Louis, Missouri, U.S.A.) was dissolved in double-deionised water and filtered through a 0.2 μm pore size Filtropur S filter (SARSTEDT AG & Co., Germany). The dye concentration was determined by U.V. absorbance (ε_412nm_ = 36.000 M^−1^cm^−1^). A 1 mM dye concentration stock solution was prepared and stored at −25 °C until use. A 10 mg/mL protein stock concentration was prepared for every experiment and diluted exactly 10X in the final solution of interest, with a 5 µM ThT concentration. A 70 μL portion of this mixture was loaded into each well in the nonbinding, clear-bottom, black 384-well plate (LOT 556276, from Nalge Nunc International - USA) and sealed using AMPLiseal Transparent Microplate Sealer (Greiner Bio-One, Frickenhausen, Germany). Each condition was present on a plate in 3 replicas, and the measurement was repeated at least twice for each condition independently using fresh material. The fluorescence measurements were performed on a PHERAstar® Plus Microplate Reader (BMG LABTECH) using a 440/480 excitation/emission window and at a controlled temperature of 30 ℃. The plates were agitated before each cycle in an orbital shaking width of 3 mm for 20 seconds. Other parameters were: positioning delay of 0.1 s; focal height was optimised for each experiment, but it was always in the range of 7.6-7.9 mm; gain of 1000; number of flashes = 100. The number of cycles and the delay time were adjusted properly to obtain the total kinetic time desired.

Turbidity measurements were performed by detecting the absorbance at 400 nm using a Greiner CELLSTAR® 384 well microplate (Sigma-Aldrich). The solution volume was kept at 70 µL at a constant temperature of 30°C. The experiments were performed in triplicate and repeated twice with fresh new sample preparation.

### 2.4 Circular Dichroism (CD)

Far UV-CD spectra (190-260 nm) were measured in a Jasco J-810 spectrometer (Jasco Corporation, Japan) equipped with a Peltier control system and a quartz cell with a 1 mm path length. The spectra were recorded from 260 to 190 nm, with a scanning speed of 50 nm/min, spectral bandwidth of 1 nm and Digital Integration Time of 2 s. All the protein stock solutions were at a minimum concentration where the dilution in the appropriated buffer solution was at least 20-fold, with a final protein concentration of 0.1 mg/mL. Thermal denaturation was performed by monitoring the ellipticity at 215 nm in the range from 20 to 66°C using a heating rate of 60°C/h. A maximum HT voltage of 600 V applied to the photomultiplier tube was used as a criterion for the data cutoff in the far-UV.

### 2.5 Negative-Stain Transmission Electron Microscopy

After the amyloid kinetic experiments, samples were taken directly from the microplates (section 2.3) and deposited for 1 min on a previously negative glow-discharged carbon-coated copper grid (Electron Microscopy Sciences, Hatfield, Pennsylvania). The grids were then washed with water and stained with phosphotungstic acid. Images were recorded on an FEI Morgagni 268 electron microscope (FEI Company, Eindhoven, Netherlands) operating at 100 keV.

### 2.6 Differential Interference Contrast (DIC) and Fluorescence Microscopies

The LLPS and droplet formation by Grh1 were visualised using an inverted system microscope IX71 equipped with a PicoQuant MT 200 module (PicoQuant®, Germany) under DIC and fluorescence mode with a 60x objective (NA = 1.2). The fluorescence excitation intervals were selected using the mirror unities: U-MNU2 BP360/370; Blue: U-MWB2 BP460/490; Green: U-MWIG3 BP530-550. The long-pass filters HQ405LP, HQ460LP, BLP01-488R, HQ550LP and HQ690/70m were used. Unless otherwise stated, the Grh1 droplets formation was induced by a 10X dilution method using a stock solution of 10 mg/mL. Solutions were mainly composed of 20 mM Na_2_HPO_4_/NaH_2_PO_4_ + X% (W/V) polyethylene glycol (PEG) 8000 (ThermoFisher) adjusted to the desired pH and with X varied from 2-12%. All the samples were incubated for 20 minutes in RT prior to measurements. The image acquisition was performed by adding 5uL drop solution in the centre of coverslips 20 x 20 mm (Deckgläser). Liquid droplet permeability was studied using 1X SYPRO™ Orange Protein Gel Stain (5,000X Concentrate in DMSO - Thermo Fisher Scientific).

### 2.7 Image, data, and statistical analysis

For the assessment of digital images captured through Differential Interference Contrast (DIC) and fluorescence microscopy, quantitative and qualitative analyses were conducted utilizing the SymPhoTime 64 software package (PicoQuant GmbH, Germany). The micrographs had their brightness and contrast adjustments meticulously carried out through Fiji histogram stretching techniques. All data and statistical analyses were performed and plotted using Origin 2022 (GraphPadSoftware).

### 2.8 Steady-state Fluorescence Spectroscopy

Fluorescence data were collected using a FluoroMax®-4 spectrofluorometer controlled by FluorEssence™ software (Horiba Jobin Yvon GmbH, Munich, Germany). For ThT fluorescence, the dye concentration was adjusted to 5 µM, and an excitation wavelength of 440 nm was used, with slits set to 2 nm.

### 2.9 Electron Paramagnetic Resonance

Grh1 was initially purified via gel filtration chromatography, using a buffer containing 5 mM 1,4-Dithiothreitol (DTT). Subsequent removal of DTT was achieved with a Hi-Trap desalting column (GE Healthcare) equilibrated in buffer A without β-mercaptoethanol. The eluted Grh1 was immediately incubated with a 10-fold molar excess of the spin-label reagent 1-oxyl-3-methanesulfonylthiomethy-2,5-dihydro-2,2,5,5-tetramethyl-1H-pyrrole (MTSL), and allowed to react overnight at 4 °C. Residual MTSL was then removed using the same desalting column and buffer solution, followed by protein concentration via Amicon Ultra 10 MWCO (Millipore). EPR Spectroscopy of Spin-Labeled Grh1: The EPR spectra of the spin-labeled Grh1 with varied concentrations of unlabelled ACBP were recorded using a JEOL FA-200 X-band spectrometer (JEOL Ltd., Tokyo, Japan) and a microwave power setting of 10 mW, conducted at room temperature. For these measurements, samples were placed in glass capillary tubes (1.5 ID X 1.8 ID; VitroCom, Inc., NJ). The analyses were carried out using the Origin 2022 software.

## Acknowledgements

The authors thank the Fundação de Amparo à Pesquisa do Estado de São Paulo (FAPESP) (Grants No. 2022/06006-0, 2021/10465-7, 2009/54044-3, 2015/50366-7, and 2020/15542-7), Conselho Nacional de Desenvolvimento Científico e Tecnológico (CNPq) (Grant no. 306682/2018-4) and Coordenação de Aperfeiçoamento de Pessoal de Nível Superior (CAPES) for the financial support.

## Supplementary Materials

This PDF file includes:

Figs. S1 to S10

Legends for movies S1 to S2

