## Supplementary data for "The potential role of liquid-liquid phase separation in the cellular fate of the compartments for unconventional protein secretion"

### Grh1 under native (pH 7.4) and "starvation" (pH 5.4) conditions

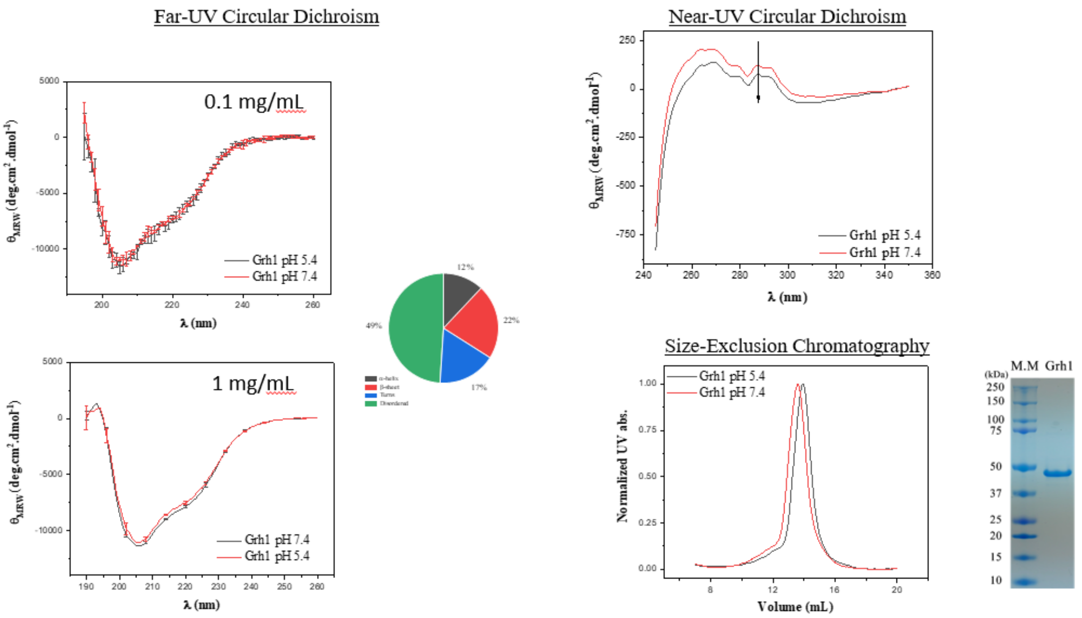


**Figure S1**: Biophysical characterisation of Grh1 at Native and Starvation pH conditions. This figure delineates the biophysical properties of Grh1, investigated through circular dichroism (CD) spectroscopy to probe the protein's secondary structure under native conditions (NC – pH 7.4) and during starvation (pH 5.4). CD spectra were recorded in the far-UV range at two protein concentrations, 0.1 and 1 mg/mL, using cuvettes with pathlengths of 0.1 and 0.01 cm, respectively. The protein concentration was maintained at 2 mg/mL with a cuvette path length of 0.1 cm for the near-UV CD measurements. Additionaly, size-exclusion chromatography (SEC) was employed to analyse the quaternary structure and oligomeric state of Grh1 under these pH conditions. An SDS-PAGE of purified Grh1 is presented.

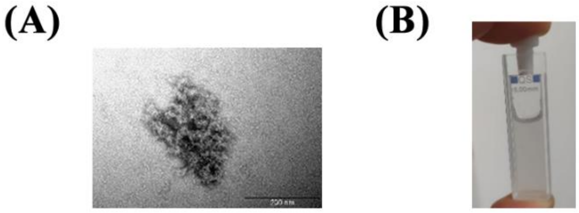

**Fig. S2:** Characterisation of Grh1 amorphous aggregation post-CD thermal denaturation in starvation-mimicking environments: (A) Electron Microscopy image displaying a typical amorphous aggregate formed by Grh1. (B) Circular Dichroism quartz cuvette post-thermal denaturation, highlighting the turbidity of the solution indicative of aggregation.

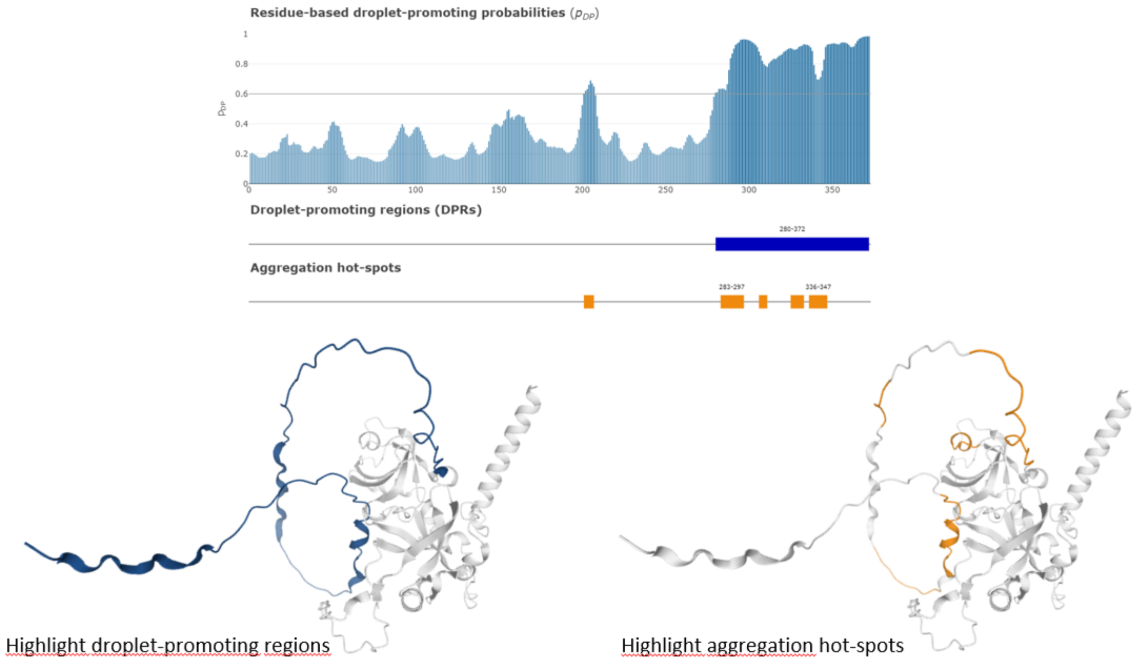


**Figure S3:** Disorder, Amyloidogenic of droplet-promoting region predictions in Grh1 using FuzPred. The regions with a high tendency for droplet promotion (blue) and amyloid formation (orange) are highlighted in the Alphafold model of Grh1.

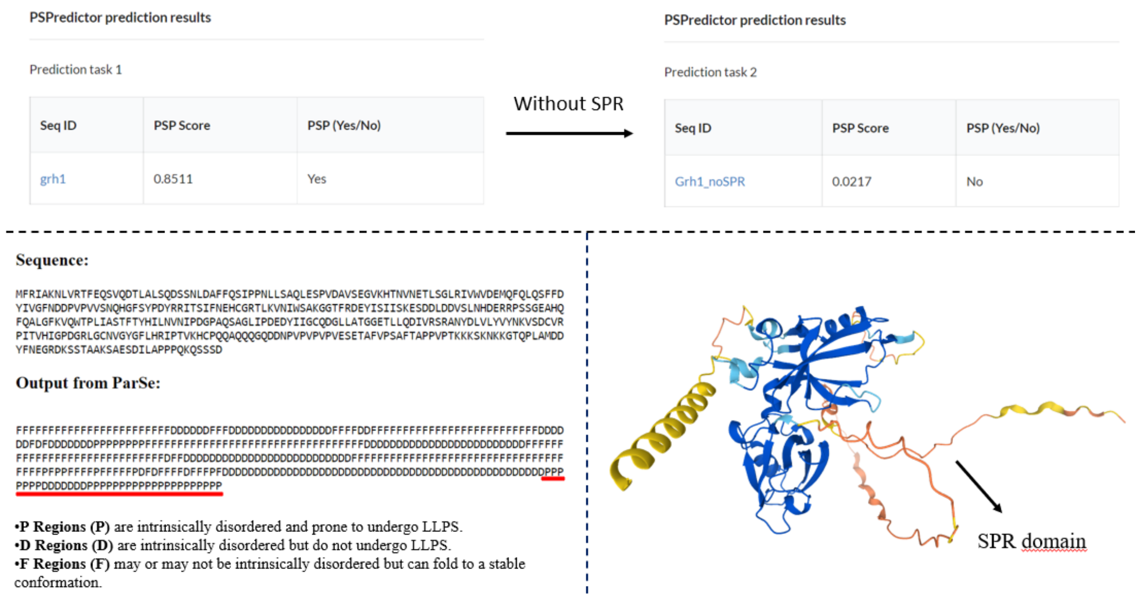


**Figure S4:** Prediction of regions involved in LLPS using the Grh1 sequence and two separate predictors: PSPredictor and ParSe. The analyses explored the high propensity for LLPS in Grh1 and mapped it to the intrinsically disordered C-terminal (Serine and Proline-Rich domain, or just SPR domain).

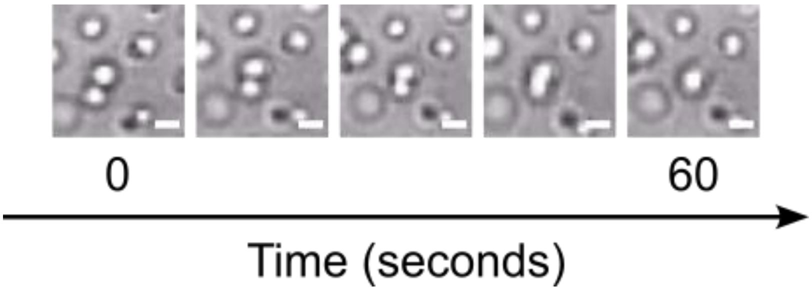


**Figure S5:** Representative fusion event of Grh1 droplets monitored by DIC microscopy. The protein is in starvation conditions. Scale bar = 1 μm.

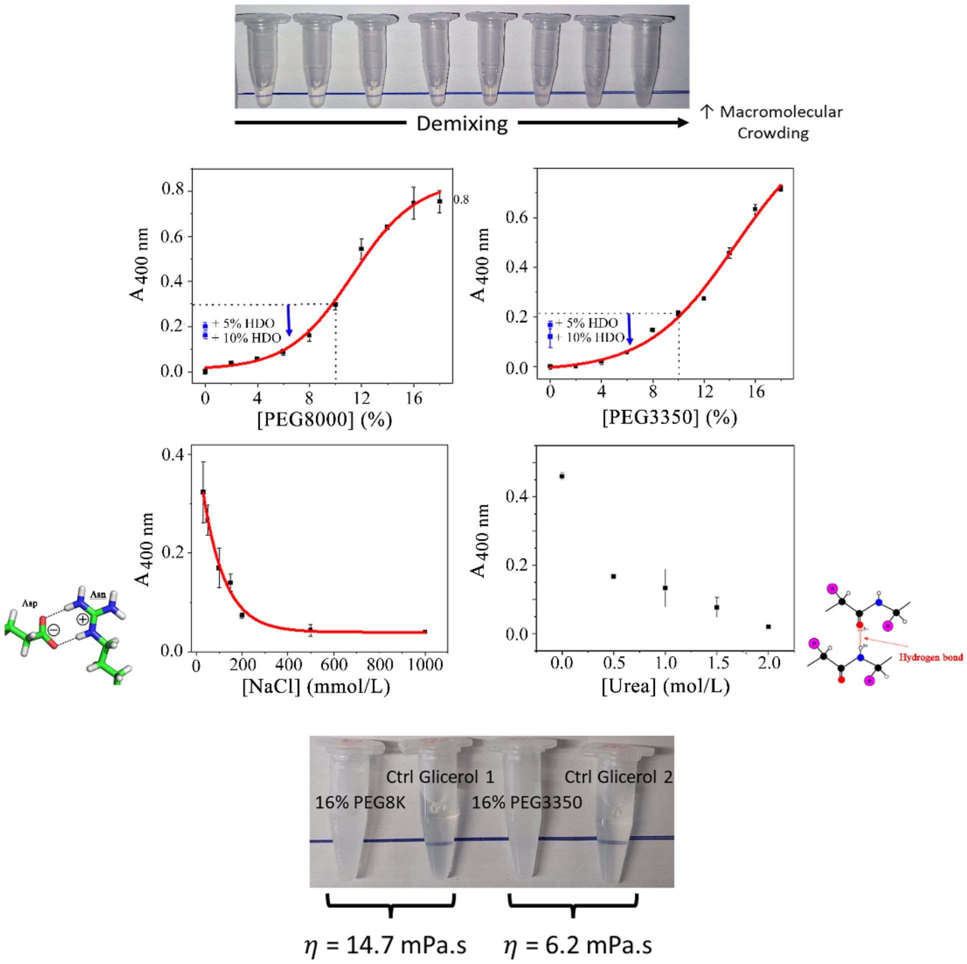


**Figure S6:** Physicochemical properties of Grh1 in its droplet state. The figure illustrates that the formation of Grh1 droplets is unaffected by the type of crowding agent used (PEG3350 and PEG8000 are presented here). It also demonstrates the sensitivity of these droplets formed in either 10% PEG8000 or PEG3350 to various de-stabilising agents: 5% and 10% of 1.6 Hexanediol (named HDO in blue; affecting weak hydrophobic interactions), NaCl (affecting electrostatic interactions), and Urea (affecting hydrogen-bound formation), at concentrations lower than those required for protein denaturation as referenced in [5]. Furthermore, comparative analysis with glycerol is presented to discern the specific contribution of macromolecular crowding, as opposed to increased solution viscosity, to the droplet formation observed in the presence of polyethylene glycol (PEG). This distinction is visually depicted in the accompanying bottle image. For the preparation of the control samples with glycerol, the conditions to reach each desired viscosity were taken from:

http://www.met.reading.ac.uk/~sws04cdw/viscosity_calc.html (accessed in 2022)

The red curves represent the fit using a Boltzmann sigmoidal function for conditions with PEG and an exponential decay model for the NaCl titration. These fits are not chosen to represent specific fitting models but to enhance the visualisation of the overall trends in the data. Detailed exploration of models accounting for these variations will be addressed in future studies.

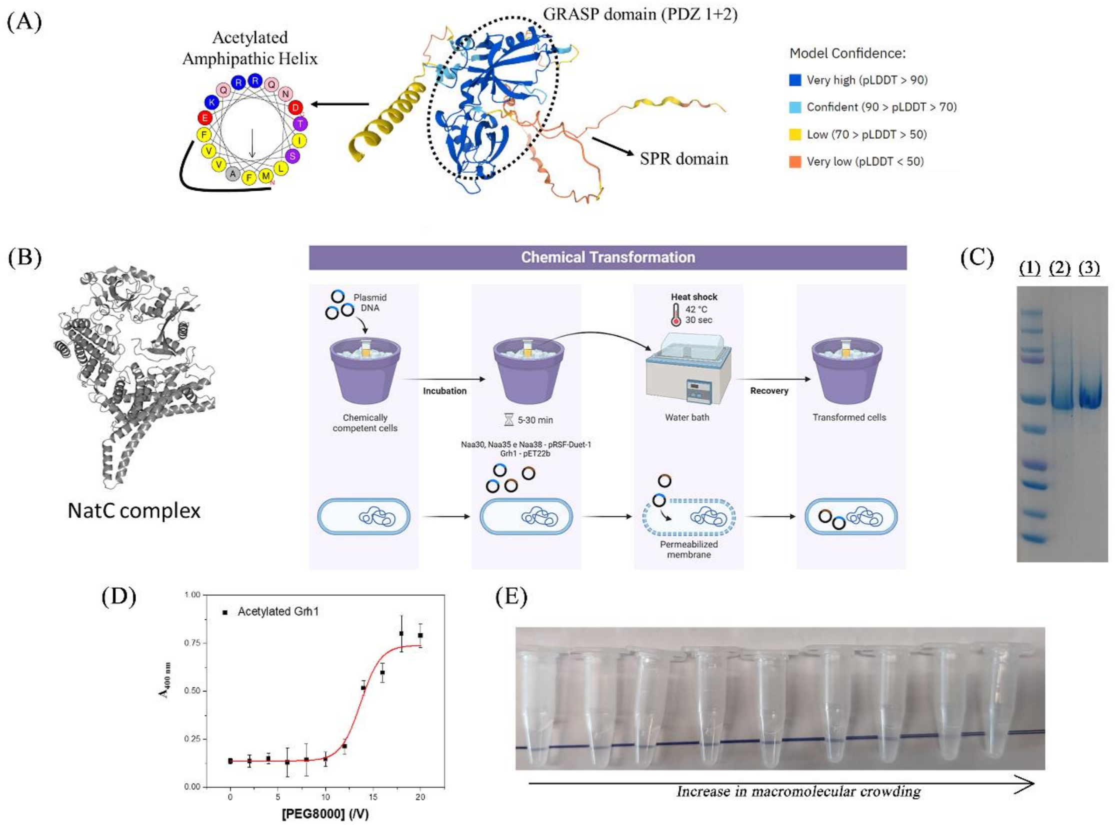


**Figure S7:** *In vivo* Acetylation of Grh1 mediated by the NatC Complex. (A) The structural model of Grh1 was generated by AlphaFold and annotated to denote the different domain regions. The model is colour-coded based on the per-residue confidence score (predicted Local Distance Difference Test, pLDDT). Notably, it includes a predicted amphipathic α-helix at the N-terminus. (B) Schematic of the co-transformation protocol employed in *E. coli*, illustrating the simultaneous introduction of the NatC complex subunits (Naa30, Naa35, and Naa38) harboured in pRSF-duet 1 (KanR), along with the Grh1 construct in pET22b (AmpR). (C) SDS-PAGE analysis displaying the bands corresponding to purified Grh1 (lane 2) and acetylated Grh1 (lane 3). Lane 1 serves as the molecular weight standard. (D) Turbidity assay results of acetylated Grh1, presented as a function of the concentration of the crowding agent PEG8000. The assay's absorbance was measured at 400 nm. (E) Graphical depiction of the turbidity data corresponding to the assay described in (D), providing a visual interpretation of the impact of acetylation on Grh1 condensation under macromolecular crowding conditions induced by PEG8000.

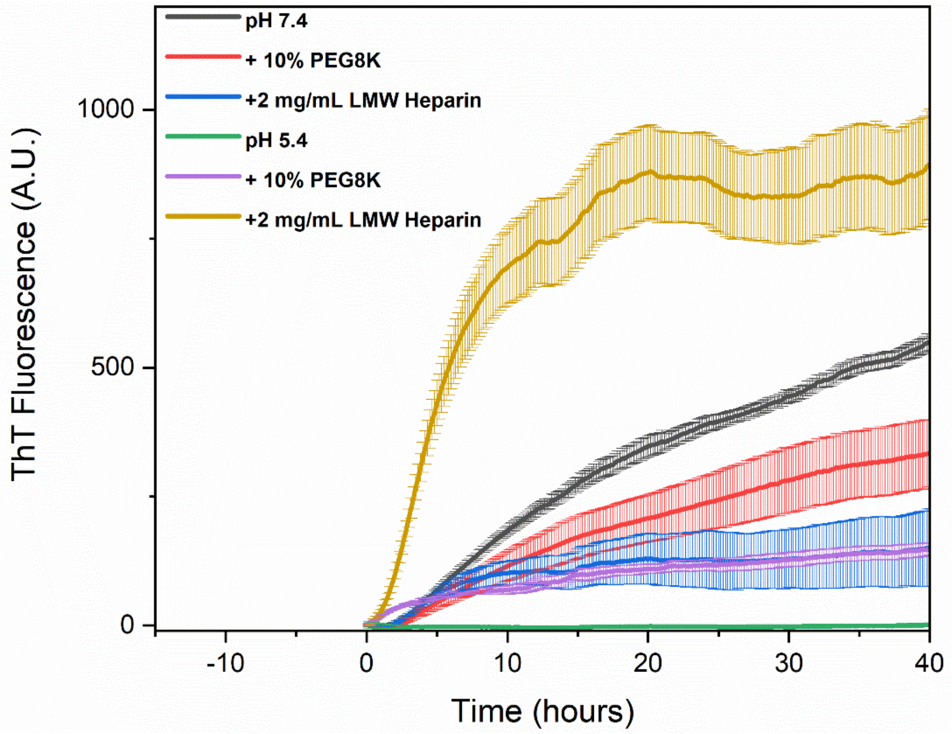


**Figure S8:** Kinetics of amyloid formation in Acetylated Grh1 assessed via Thioflavin T (ThT) Fluorescence. This set of experiments tracks the amyloidogenesis of acetylated Grh1 by measuring the fluorescence intensity of ThT, a dye that binds to amyloid fibrils. The assays were conducted at physiological pH 7.4 and acidic pH 5.4, which simulate normal cellular conditions (NC) and starvation-induced environments. The influence of low molecular weight (LMW) heparin at a concentration of 2 mg/mL and 10% polyethylene glycol (PEG) 8000 on amyloid formation was investigated.

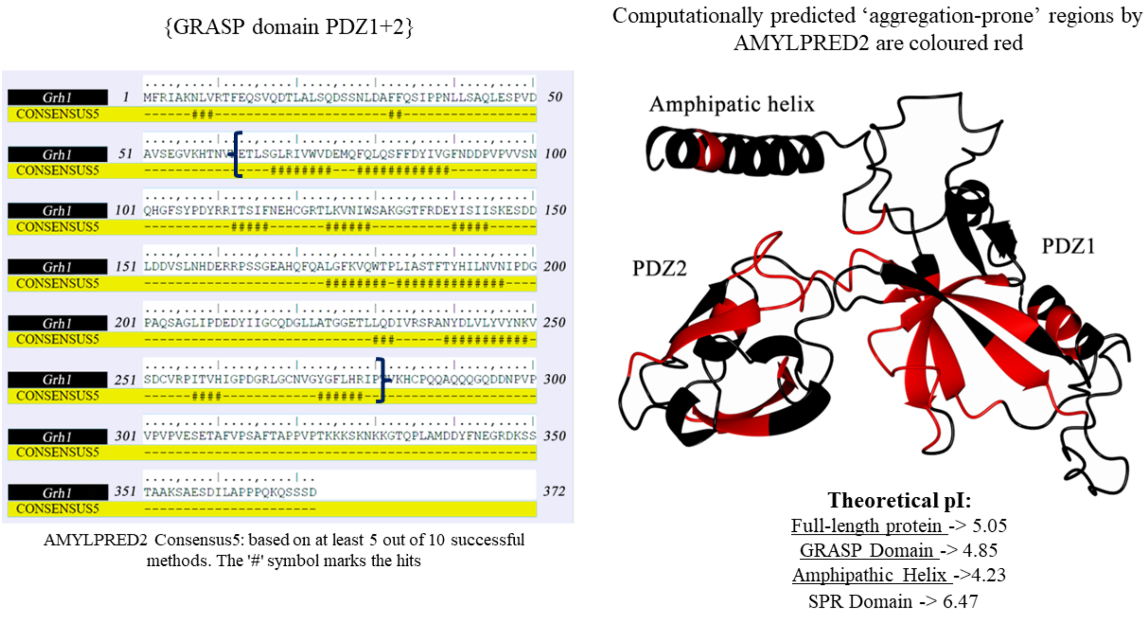


**Figure S9:** Identification of aggregation-prone regions in Grh1 utilising Amylpred2. This figure represents the consensus prediction of amyloidogenic sequences within Grh1 as determined by Amylpred2. The regions receiving concordant predictions from more than five independent algorithms are denoted with hash marks (#). The extent of the GRASP domain, encompassing PDZ1 and PDZ2, is demarcated by braces ({}). On the structural model of Grh1, these Amylpred2-identified regions are accentuated in red for enhanced visibility, with the SPR domain omitted to simplify the representation. Additionally, the distinct subunits' isoelectric points (pIs) were computationally estimated using the ProtParam tool.

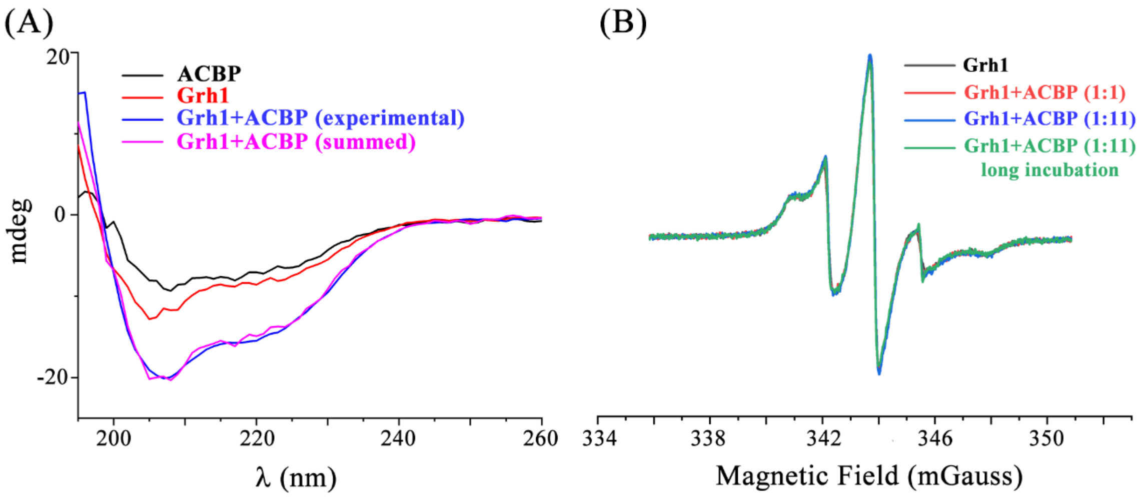


**Figure S10:** Exploring Grh1-ACBP Interactions in Soluble Conditions. We conducted experiments to study the interaction between Grh1 and ACBP under soluble conditions, employing Circular Dichroism (CD) and Electron Spin Resonance (ESR) spectroscopy. Both techniques, represented in panels (A) for CD and (B) for ESR, were performed at room temperature using Buffer A to ensure consistent environmental conditions for the interactions. For experimental details, please refer to the Material and Methods section.

**Movie S1:** Video of Grh1 droplets in starvation conditions using DIC microscopy.

**Movie S2:** Video of Grh1 droplets in starvation conditions using DIC microscopy. The video exemplifies one representative fusion event present in the solution that happens over time.
